# Quantification of genetic components of population differentiation in UK Biobank traits reveals signals of polygenic selection

**DOI:** 10.1101/357483

**Authors:** Xuanyao Liu, Po-Ru Loh, Luke J. O’Connor, Steven Gazal, Armin Schoech, Robert M. Maier, Nick Patterson, Alkes L. Price

## Abstract

The genetic architecture of most human complex traits is highly polygenic, motivating efforts to detect polygenic selection involving a large number of loci. In contrast to previous work relying on top GWAS loci, we developed a method that uses genome-wide association statistics and linkage disequilibrium patterns to estimate the genome-wide genetic component of population differentiation of a complex trait along a continuous gradient, enabling powerful inference of polygenic selection. We analyzed 43 UK Biobank traits and focused on PC1 and North-South and East-West birth coordinates across 337K unrelated British-ancestry samples, for which our method produced close to unbiased estimates of genetic components of population differentiation and high power to detect polygenic selection in simulations across different trait architectures. For PC1, we identified signals of polygenic selection for height (74.5±16.7% of 9.3% total correlation with PC1 attributable to genome-wide genetic effects; P = 8.4×10^−6^) and red hair pigmentation (95.9±24.7% of total correlation with PC1 attributable to genome-wide genetic effects; P = 1.1×10^−4^); the bulk of the signal remained when removing genome-wide significant loci, even though red hair pigmentation includes loci of large effect. We also detected polygenic selection for height, systolic blood pressure, BMI and basal metabolic rate along North-South birth coordinate, and height and systolic blood pressure along East-West birth coordinate. Our method detects polygenic selection in modern human populations with very subtle population structure and elucidates the relative contributions of genetic and non-genetic components of trait population differences.

## Introduction

The genetic architecture of human complex traits is highly polygenic^1–7^. Natural selection on polygenic traits could occur in polygenic fashion, via small shifts in allele frequencies across a large number of loci^8,9^. Signals of polygenic selection have been detected for several complex traits, including height and body mass index (BMI), by correlating SNP effect sizes from genome-wide association studies (GWAS) with population differences in allele frequencies or by computing singleton density scores^10–17^. However, methods for detecting polygenic selection generally restrict their analyses to genome-wide significant associated SNPs or relatively small sets of top associated SNPs, which generally capture only a small proportion of trait heritability^18^. This may limit power to detect polygenic selection for highly polygenic traits and precludes estimation of the genetic component of population differences in phenotype, a fundamental population genetic quantity.

In this study, we developed a method, PopDiff, that quantifies the genetic component of population differences in phenotype, using association statistics from genome-wide SNPs and linkage disequilibrium (LD) between SNPs; the method can be applied to continuous gradients of genetic ancestry, which are often an effective way to model subtle population structure^19–21^. A significantly non-zero value of the genetic component after accounting for effects of genetic drift indicates a signal of polygenic selection. We applied PopDiff to 43 UK Biobank traits (*N*=337K unrelated British-ancestry samples^22^), analyzing three continuous gradients: the top principal component (PC1) and North-South and East-West birth coordinates. We detected signals of polygenic selection for several traits, including traits not previously reported to be under polygenic selection.

## Results

### Overview of methods

Population differences in phenotype may have genetic and/or environmental components. Our method, PopDiff, quantifies the genetic component of population differences in phenotype to detect signals of polygenic selection. We first consider the special case of two discrete subpopulations and no LD. In this case, given a phenotype whose mean differs between the two subpopulations, an unbiased estimate of the genetic component of the phenotypic difference can be obtained by summing the estimated contribution of each SNP to the phenotypic difference, based on the product of the difference in allele frequency and the estimated effect size. (Throughout this paper, “genetic component” is defined as the component explained by a specified set of SNPs and may exclude other genetic effects.) Although genetic differences between subpopulations can arise due to either genetic drift or polygenic selection, the effects of genetic drift vary stochastically across the genome, such that standard errors computed using a block-jackknife include the effects of genetic drift and enable a statistical test for polygenic selection. It is straightforward to generalize from two discrete subpopulations to a pre-specified continuous gradient of genetic ancestry (e.g. based on principal components (PCs) or birth coordinates) by replacing the difference between subpopulations with the correlation to the continuous gradient. In the presence of LD, which can cause effects of linked SNPs to be double-counted, we multiply estimated SNP effects by the inverse of the LD matrix to correct for LD^23^. To reduce noise, we regularize the LD matrix as described previously^24^. (This regularization can introduce a conservative bias in estimates but increases power to detect polygenic selection; we thoroughly investigate this bias-variance tradeoff in our simulations). We note that analyses of polygenic selection can potentially be confounded by uncorrected population stratification^28,29^. We correct for 10 PCs when estimating marginal SNP effects, but careful consideration of possible uncorrected population stratification is warranted (see Population stratification section).

In detail, we estimate the genetic component Δ*G* of population differentiation Δ*Y* (defined as the correlation between continuous ancestry gradient and phenotype *Y*) via

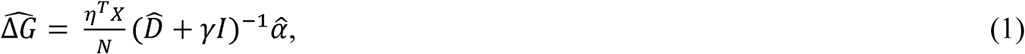

where *η* is a pre-specified *N* x 1 vector quantifying a continuous gradient of genetic ancestry across samples (normalized to mean 0 and variance 1), *X* is the *N* x *M* matrix of normalized genotypes, 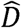 is an *M* x *M* banded LD matrix computed using all samples, *γ* is a scalar regularization parameter^24^, *I* is the *M* x *M* identity matrix, 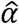 is the *M* x 1 vector of normalized estimated marginal (inclusive of LD) effect sizes for each SNP, *M* is the number of SNPs, and *N* is the number of samples. We note that 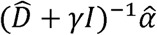 is an estimate of normalized *causal* SNP effect sizes *β*, so that 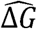 is an estimate of 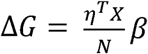. The genetic proportion of population differentiation, denoted %*G*, is estimated as 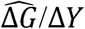. As noted above, we estimate the standard error of 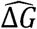 using a block-jackknife that includes the effects of genetic drift, so that a significantly nonzero 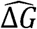 indicates a signal of polygenic selection. Details of our PopDiff method are described in the Methods section; we have released open-source software implementing the method (see URLs).

In this study, we analyzed data from 337,536 unrelated British-ancestry samples from the full UK Biobank release^22^ (see URLs). We considered 43 highly heritable traits and three continuous ancestry gradients *η*: PC1 of the 337,536 unrelated British-ancestry samples, North-South birth coordinates and East-West birth coordinates. (We did not consider lower PCs, which correspond to exceedingly subtle genetic effects: *F*_ST_ <0.0001; Table S2 of ref. 21). We applied our PopDiff method to 67 (*η*, trait) pairs for which the population differentiation Δ*Y* (correlation between *η* and phenotype *Y*) had absolute value greater than 0.01 (Table S1). In all analyses, we computed the LD matrix 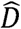 using the full set of 337,536 samples. In analyses of North-South and East-West birth coordinates, we computed marginal effect size estimates 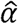 using the full set of 337,536 samples, correcting for 10 PCs. In analyses of PC1, this choice of 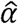 would be mathematically guaranteed to produce a 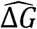 estimate of 0 (since correlations to PC1 are subtracted out, see Methods), and for this reason we instead computed 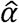 using 10 random (non-overlapping) subsets of 33,754 samples (correcting for 10 PCs within each subset) and meta-analyzed the results.

### Simulations

We performed simulations using real UK Biobank genotypes to assess the bias, type I error and power of our PopDiff method. We performed simulations using both PC1 and North-South birth coordinates. We used unrelated British-ancestry samples (*N*=337,536) and genome-wide genotyped SNPs (*M*=516,086 SNPs after QC; see Methods); our simulations used exactly the same sample set and SNP set as our analyses of real traits. Phenotypes were simulated using an additive model with SNP-heritability set to 0.2, similar to most UK Biobank traits^25,26^ (see Methods). The proportion of causal SNPs was set to 100% (the default value), 10% or 1%, and the regularization parameter *γ* was set to 0.1 (as in ref. 24), 0.2, 0.5 or 1.0. In PC1 simulations, we estimated 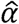 using a random subset of unrelated British-ancestry samples (*N*=33,754, correcting for 10 PCs within the subset), analogous to our analyses of real traits.

We performed null simulations (heritable phenotype with Δ*Y*= 0.11, *ΔG*= 0, %*G* = 0) to assess bias and type I error. We first performed simulations using PC1. Estimates of Δ*G* and *%G* = Δ*G*/Δ*Y* were unbiased at all values of *γ*, although estimates were extremely noisy at *γ*=0.1 (Figure 1A and Table S2A). Type I error was properly controlled at all values of *γ* (Figure 1B and Table S2B; conservative at *γ*=0.1 due to noisy estimates), and jackknife standard errors (s.e.) were similar to empirical standard deviations (s.d.) of Δ*G* estimates (Table S2A). We also performed simulations using North-South birth coordinates. Once again, estimates of Δ*G* and %*G*= Δ*G*/Δ*Y* were unbiased at all values of *γ*, although estimates were extremely noisy at *γ*=0.1 (Figure S1A and Table S3A). Type I error was properly controlled for *γ*=0.1, 0.2, 0.5 but not for *γ* = 1.0 (Figure S1B and Table S3B; conservative at *γ* = 0.1 due to noisy estimates), and jackknife s.e. were similar to empirical s.d. for all values of *γ* (Table S3A); we chose *γ*=0.5 as the default setting (see below). Notably, we obtained similar results at different values of the proportion of causal SNPs (100%, 10% or 1%), both for simulations using PC1 (Table S4) and for simulations using North-South birth coordinates (Table S5).

**Figure 1.**
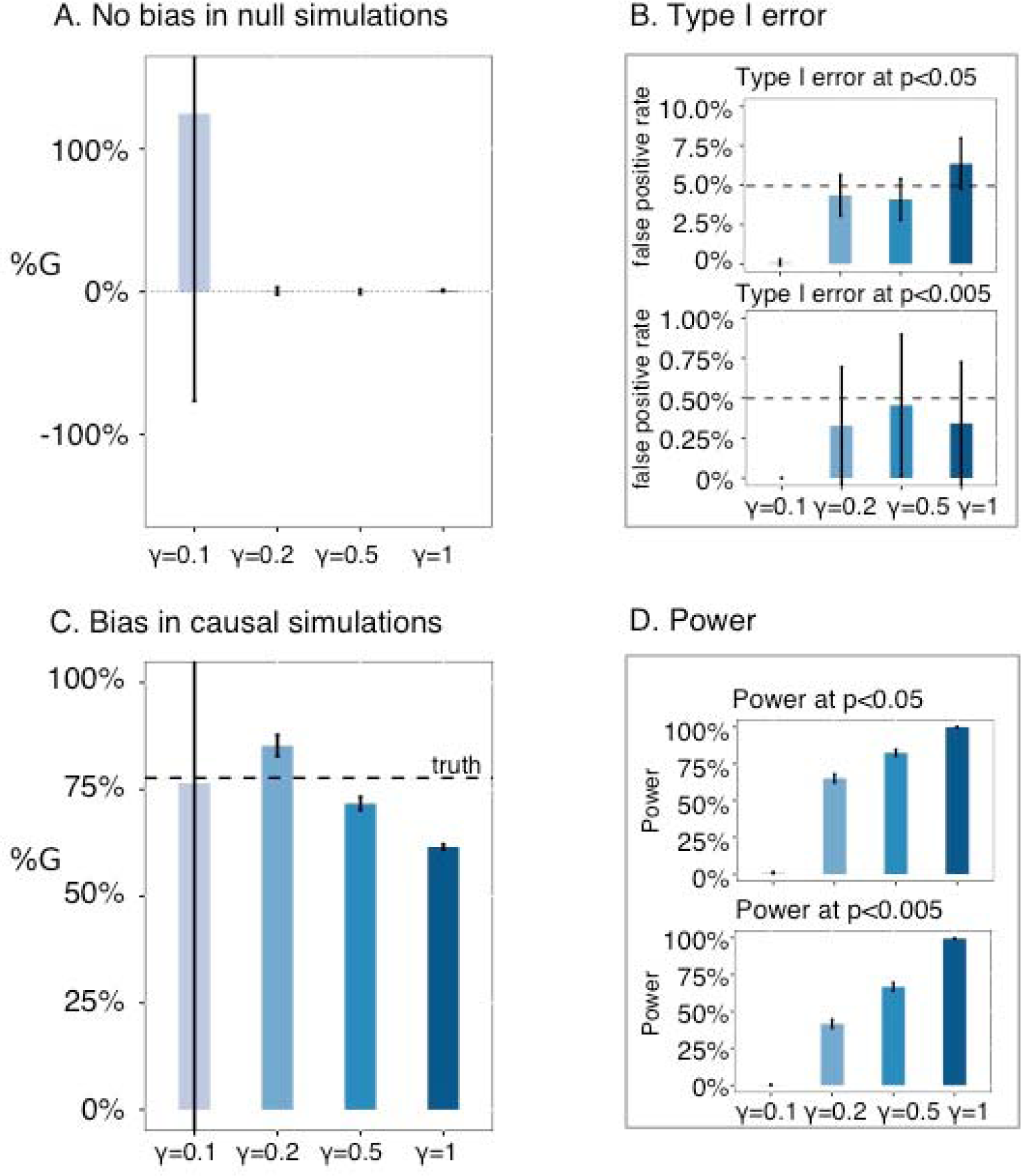
Null and causal PC1 simulations to evaluate bias, type I error and power. (A) No bias in null simulations. We report the bias in %*G* estimates for different values of γ. (B) Type I error. We report type I error at p<0.05 and type I error at p<0.005 for different values of γ. (C) Bias in causal simulations. We report the bias in *%G* estimates for different values of γ. (D) Power. We report power at p<0.05 and power at p<0.005 for different values of γ. Error bars represent 95% confidence intervals. Numerical results are reported in Table S2 and Table S6.

We performed causal simulations (Δ*Y*= 0.11, Δ*G* = 0.086, %*G* = 78%) to assess bias and power. We first performed simulations using PC1. Estimates of Δ*G* and *%G*= Δ*G*/Δ*Y* were unbiased but extremely noisy at γ=0.1, slightly upward biased at γ =0.2, and slightly conservative at γ =0.5 and γ =1.0 (Figure 1C and Table S6A). Power increased as a function of y, with high power at γ =0.5 and very high power at γ =1.0 (Figure 1D and Table S6B). We also performed simulations using North-South birth coordinates. Estimates of Δ*G* and *%G*= Δ*G*/Δ*Y* were upward biased and extremely noisy at γ =0.1, slightly upward biased at γ =0.2, close to unbiased at γ =0.5, and conservative at y=1.0 (Figure S1C and Table S7A). Power increased as a function of y, with very high power at γ =0.5 and y=1.0 (Figure S1D and Table S7B). As in null simulations, we obtained similar results at different values of the proportion of causal SNPs (100%, 10% or 1%), both for simulations using PC1 (Table S8) and for simulations using North-South birth coordinates (Table S9). Based on the results of both null and causal simulations, we chose γ =0.5 as the default regularization parameter value in all of our analyses of real traits, as this parameter value consistently controls false positives, produces close to unbiased or slightly conservative estimates, and achieves high power. (For completeness, we also report results of secondary analyses at γ =1 in our analyses of real traits.)

### Polygenic selection along PC1 in UK Biobank

We considered 43 UK Biobank traits, restricting to 337,536 unrelated British-ancestry samples (average *N* = 321,389 phenotyped samples; Table S10). We defined a continuous ancestry gradient *η* based on PC1 of the full set of 337,536 samples, representing a north-south axis separating southern England from Northern Ireland^21^. We applied our PopDiff method to 22 (PC1, trait) pairs for which the population differentiation Δ*Y* (correlation between PC1 and phenotype *Y*) had absolute value greater than 0.01 (Table S1). All analyses were corrected for 67 hypotheses tested, which include other choices of *η* (see below).

Results are displayed in Figure 2 and Table S11. We identified two traits with statistically significant %*G* for PC1 (*p*<0.05/67=7.5×10^−4^), implicating polygenic selection: height and red hair pigmentation. For height (Δ*Y*=0.093; individuals with ancestry from southern England are taller on average than individuals with ancestry from Northern Ireland), our estimate of %*G* was 74.5% (s.e.=16.7%; *p*=8.4×10^−6^), implying that differences in height along PC1 are primarily due to selection and cannot be explained by genetic drift. We note that height has previously been reported to be under polygenic selection^10–16^. For red hair pigmentation (Δ*Y*=−0.039; red hair is more common in individuals with ancestry from Northern Ireland than in individuals with ancestry from southern England), our estimate of %*G* was 95.9% (s.e.=24.7%; *p*=1.1×10^−4^), implying that differences in red hair pigmentation along PC1 are primarily due to selection and cannot be explained by genetic drift. We note that the genetic architecture of red hair pigmentation includes large-effect loci, with 12 genome-wide significant loci explaining 7.2% of trait variance (Table S12, comparable to 6.9% in ref. 27). We repeated our analysis after removing these 12 loci and surrounding regions (±1Mb), and confirmed that the signal of polygenic selection remained (%*G*=73.2%, s.e.=21.3%; *p*=6.1×10^−4^). This demonstrates that polygenic selection can affect traits whose genetic architectures include large-effect loci. We are not currently aware of previous evidence of polygenic selection on red hair pigmentation, although a previous study reported polygenic selection on skin pigmentation^11^. In secondary analyses at γ=1, %*G* estimates for height and red hair pigmentation were lower (consistent with Figure 1C), but remained statistically significant (Table S13A).

**Figure 2.**
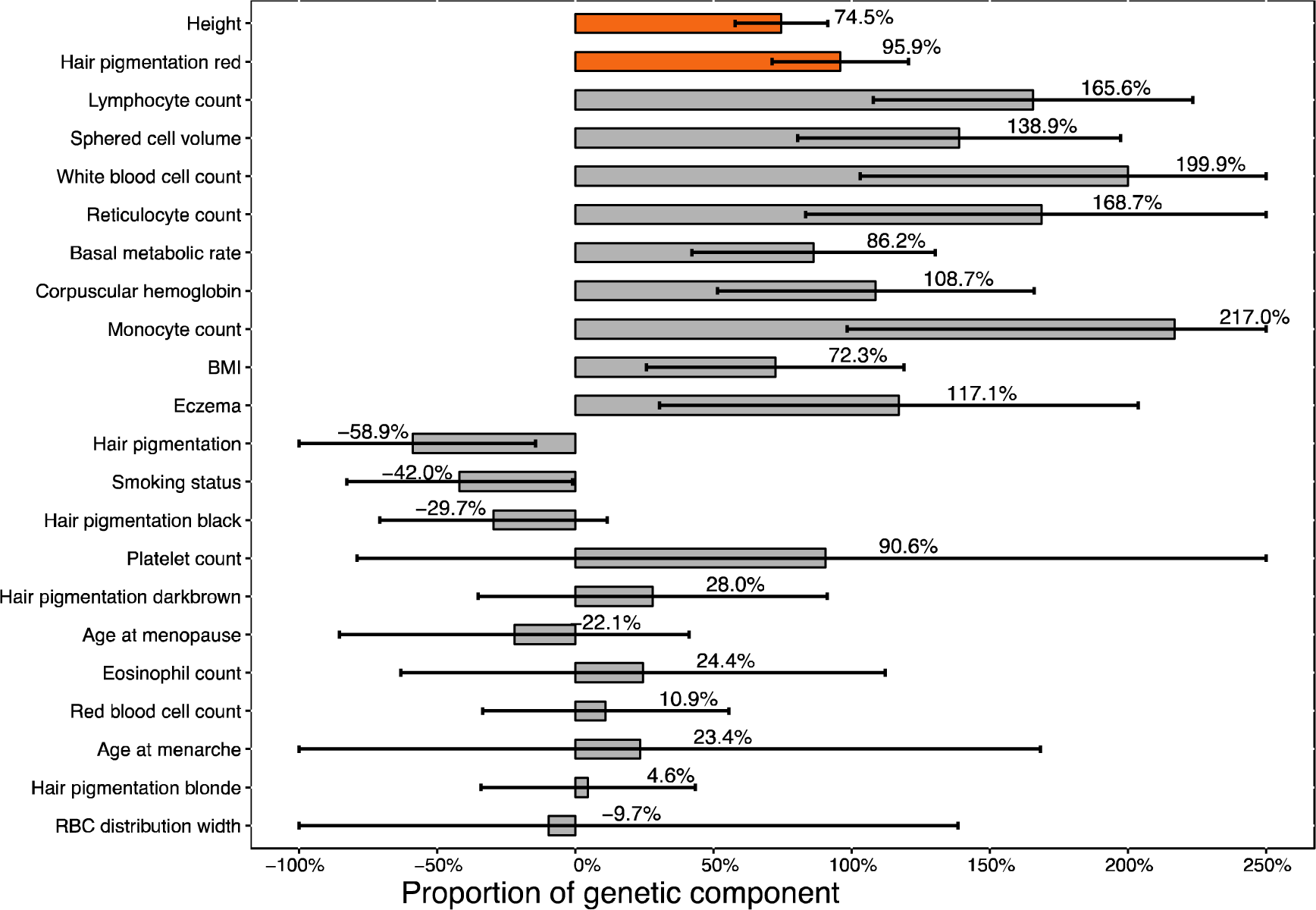
Estimates of genetic components of population differentiation (%*G*) and inference of polygenic selection along PC1 in UK Biobank. We report point estimates and standard errors for %*G* along PC1 for 22 UK Biobank traits for which Δ*Y* (correlation between PC1 and phenotype *Y)* had absolute value greater than 0.01. Traits are ranked by statistical significance of nonzero %*G*. Traits with Bonferroni-significant nonzero %*G* (p < 0.05/67), indicative of polygenic section, are denoted via orange bars. Numerical results are reported in Table S11.

### Population stratification

Recent work has suggested that previous studies of polygenic selection may be confounded by uncorrected population stratification, compromising their results^28,29^. Correcting for population stratification is clearly very important in analyses of polygenic selection, as we determined that repeating our analyses with no correction for population stratification produced unstable results (Table S14; e.g. height %*G* > 2000%, *p*<10^−200^). We note that uncorrected population stratification may be either environmentally driven (driven by environmental components of population differences in phenotype) or genetically driven (driven by genetic components of population differences in phenotype, caused by polygenic selection). We are primarily concerned about the former case, as the latter case represents true-positive (not false-positive) signals of polygenic selection— although estimates of %*G* could still be inflated in the latter case.

We performed a series of analyses to assess whether our results are robust to uncorrected population stratification. We first considered simulations. Our null PC1 simulations described above (heritable phenotype with Δ*Y*= 0.11, Δ*G* = 0, %*G*= 0; Figure 1A,B), which correspond to the case of environmentally driven population stratification, achieved correct calibration. However, it is also of interest to check whether environmentally driven population stratification along other genetic gradients could lead to false-positive signals of polygenic selection along PCI. To assess this, we performed additional simulations in which we simulated environmentally driven population stratification along North-South birth coordinate (analogous to Figure S1A,B) but evaluated evidence of polygenic selection along PCI. Results are reported in Figure S2 and Table S15. We confirmed that type I error was properly.

We next considered analyses of UK Biobank traits. We repeated our PC1 analyses by estimating *a* using 10 (non-overlapping) subsets of 33,754 samples ordered by PC1 values (similar values of PC1 within each subset, so as to minimize stratification), correcting for PCs within each subset and meta-analyzing the results. We confirmed that %*G* remained statistically significant for both height (%*G*=123.0%, s.e.=14.7%; *p*=4.9×10^−17^) and red hair pigmentation (%*G*=93.54%, s.e.=35.26%; *p*=7.9×10^−3^); results for all 22 traits are reported in Table S16. We also repeated our PC1 analysis of height using family-based effect size estimates from ref. 28, and determined that the %*G* estimate along British PC1 was 95.5% (s.e.=40.8%, *p*=0.019); in secondary analyses at γ=1, the %*G* estimate remained positive, but was smaller and non-significant (Table S13B). We note that our results involving British PC1 are orthogonal to previous findings involving European PC1^28^: we repeated our analysis using a European PC1 computed using all *N*=460K European-ancestry samples^22,26^, and determined that the correlation between European PC1 and British PC1 loadings was only −0.017, and that the %*G* for height along European PC1 (relative to Δ*Y*= −2.5%) was −20.7% (s.e.=10.2%, *p*=0.043), which is consistent with the ref. 28 finding that the genetic component of the population difference in height along European PC1 has the opposite sign of the total population difference. Overall, these secondary analyses support our findings (as well as those of ref. 28).

### Polygenic selection along North-South birth coordinate in UK Biobank

We next defined a continuous ancestry gradient *η* based on North-South birth coordinate. We applied our PopDiff method to 24 (North-South birth coordinate, trait) pairs for which the population differentiation Δ*Y* (correlation between North-South birth coordinate and phenotype *Y*) had absolute value greater than 0.01 (Table S1). All analyses were corrected for 67 hypotheses tested.

Results are displayed in Figure 3 and Table S17. We identified four traits with statistically significant %*G* for North-South birth coordinate (*p*<0.05/67=7.5×10^−4^): height, BMI, basal metabolic rate and systolic blood pressure. For height (Δ*Y*=-0.091; individuals born in the southern UK are taller on average than individuals born in the northern UK), our estimate of %*G* was 124.6% (s.e.=15.9%; *p*=4.6×10^−15^), implying that differences in height along North-South birth coordinate are predominantly genetic and cannot be explained by genetic drift. For BMI (Δ *Y*=0.040; individuals born in the northern UK have larger BMI on average than individuals born in the southern UK), our estimate of %*G* was 128.2% (s.e. = 25.7%; p=5.8×10^−7^), implying that differences in BMI along North-South birth coordinate are predominantly genetic and cannot be explained by genetic drift. Both height and BMI have previously been reported to be under polygenic selection^10–16^. We note that our estimates of %*G* are not significantly larger than 100%; however, values of %*G* larger than 100% are possible if genetic and environmental geographic effects have opposite signs, as previously reported for BMI (ref. 12).

**Figure 3.**
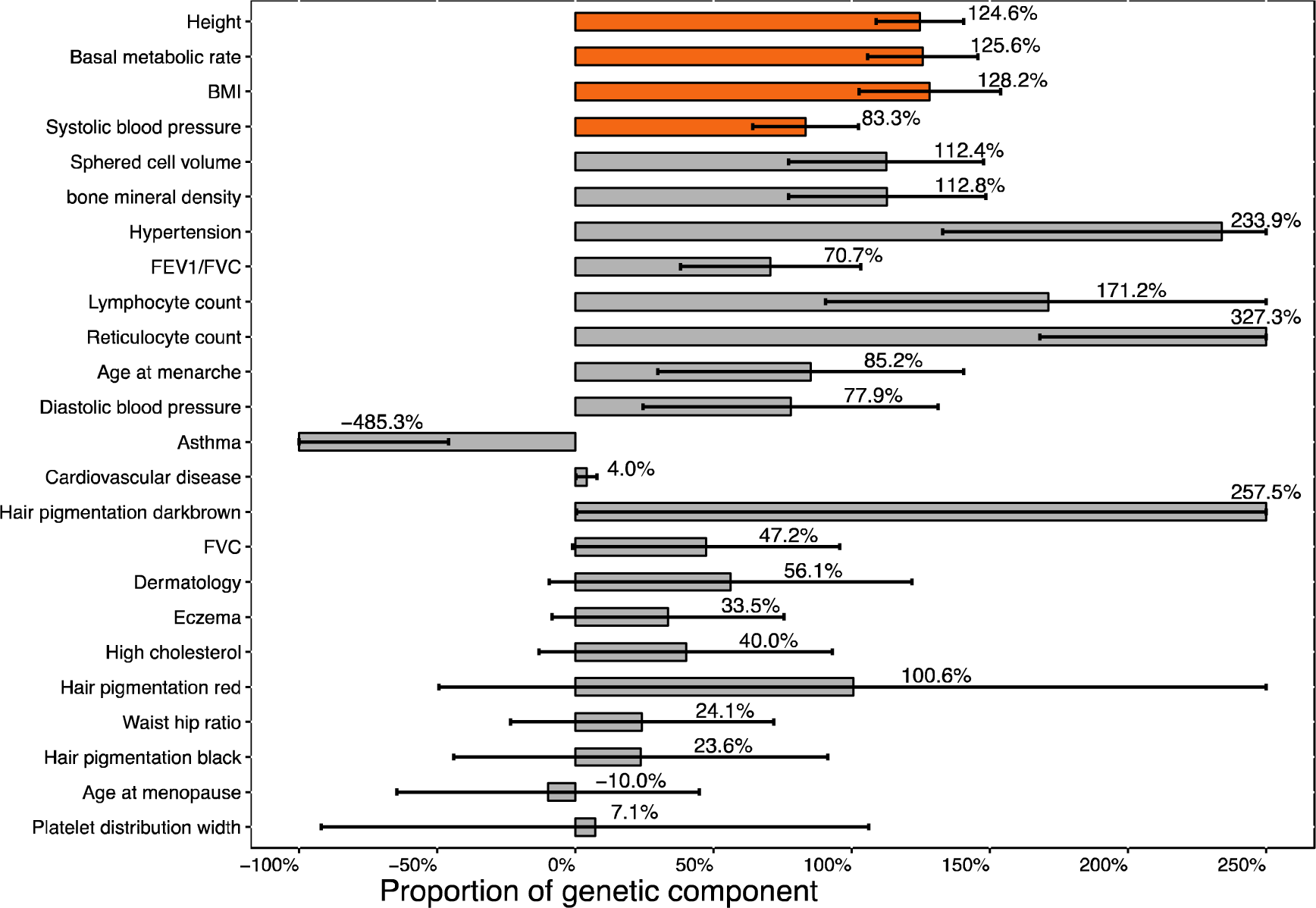
Estimates of genetic components of population differentiation (%*G*) and inference of polygenic selection along North-South birth coordinate in UK Biobank. We report point estimates and standard errors for %*G* along North-South birth coordinate for 24 UK Biobank traits for which Δ*Y* (correlation between North-South birth coordinate and phenotype *Y)* had absolute value greater than 0.01. Traits are ranked by statistical significance of nonzero %*G*. Traits with Bonferroni-significant nonzero %*G* (p < 0.05/67), indicative of polygenic section, are denoted via orange bars. Numerical results are reported in Table S17.

We are not currently aware of previous evidence of polygenic selection on systolic blood pressure and basal metabolic rate. For systolic blood pressure (Δ*Y*=0.032; individuals born in the northern UK have higher systolic blood pressure on average than individuals born in the southern UK), our estimate of %*G* was 83.3% (s.e.=19.1%; *p*=1.3×10^−5^), implying that differences in systolic blood pressure along North-South birth coordinate are primarily due to selection and cannot be explained by genetic drift. For basal metabolic rate (Δ*Y*=-0.032; individuals born in the southern UK have higher basal metabolic rate on average than individuals born in the northern UK), our estimate of %*G* was 125.6% (s.e.=20.0%; *p*=3.3×10^−10^), implying that differences in basal metabolic rate along North-South birth coordinate are primarily due to selection and cannot be explained by genetic drift. We note that both systolic blood pressure and basal metabolic rate have significant genetic correlation with height and BMI in UK Biobank data (Table S18; estimated using cross-trait LD score regression^30^). For both systolic blood pressure and basal metabolic rate, %*G* became only nominally significant (0.05/67=7.5×10^−4^<*p*<0.05) when computing association statistics using height as a covariate (Table S19), suggesting that polygenic selection on these traits may be impacted by polygenic selection on height. On the other hand, for both of these traits, %*G* remained highly significant when computing association statistics using BMI as a covariate (Table S19), although we caution that adjusting association statistics for heritable covariates can introduce collider bias^31^. In secondary analyses at γ=1, %*G* estimates were lower for most traits, but remained statistically significant (Table S20).

### Polygenic selection along East-West birth coordinate in UK Biobank

Finally, we defined a continuous ancestry gradient based on East-West birth coordinate. We applied our PopDiff method to 21 (East-West birth coordinate, trait) pairs for which the population differentiation Δ*Y* (correlation between East-West birth coordinate and phenotype *Y*) had absolute value greater than 0.01 (Table S1). All analyses were corrected for 67 hypotheses tested.

Results are displayed in Figure 4 and Table S21. We identified two traits with statistically significant %*G* for East-West birth coordinate (*p*<0.05/67=7.5×10^−4^): height and systolic blood pressure, both of which were also statistically significant in our analysis of North-South birth coordinate. For height (Δ*Y*=0.087; individuals born in the eastern UK are taller on average than individuals born in the western UK), our estimate of %*G* was 78.0% (s.e.=21.2%; *p*=2.4×10^−4^), implying that differences in height along East-West birth coordinate are primarily due to selection and cannot be be explained by genetic drift. For systolic blood pressure (Δ*Y*=-0.031; individuals born in the western UK have higher systolic blood pressure on average than individuals born in the eastern UK), our estimate of %*G* was 85.2% (s.e.=16.4%; *p*=2.4×10^−7^), implying that differences in systolic blood pressure along East-West birth coordinate are primarily due to selection and cannot be explained by genetic drift. The %*G* for systolic blood pressure remained highly significant when computing association statistics using height as a covariate (Table S22), although we caution that adjusting association statistics for heritable covariates can introduce collider bias^31^. In secondary analyses at γ=1, %*G* estimates were lower, but remained statistically significant (Table S23).

**Figure 4.**
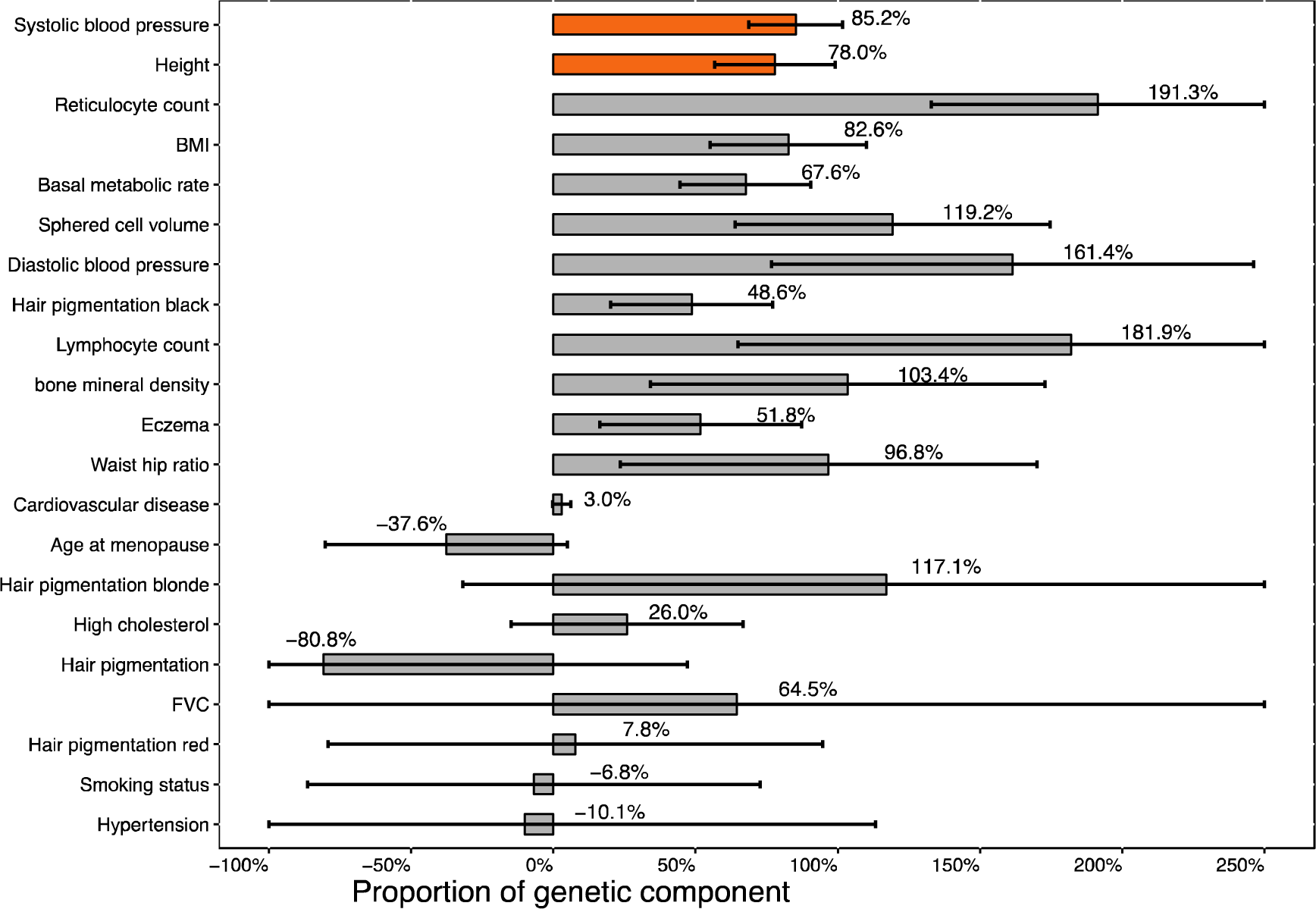
Estimates of genetic components of population differentiation (%*G*) and inference of polygenic selection along East-West birth coordinate in UK Biobank. We report point estimates and standard errors for %*G* along East-West birth coordinate for 21 UK Biobank traits for which Δ*Y* (correlation between East-West birth coordinate and phenotype *Y*) had absolute value greater than 0.01. Traits are ranked by statistical significance of nonzero %*G.* Traits with Bonferroni-significant nonzero %*G*(p < 0.05/67), indicative of polygenic section, are denoted via orange bars. Numerical results are reported in Table S21.

## Discussion

We developed a method, PopDiff, that quantifies the genetic component of population differences in phenotype to detect signals of polygenic selection. The method was well-powered in simulations and analyses of real UK Biobank traits in detecting polygenic selection within British-ancestry samples, which have very subtle population structure. We identified several traits under polygenic selection, including traits previously reported to be under polygenic selection (height and BMI) and other traits (red hair pigmentation, systolic blood pressure and basal metabolic rate).

PopDiff is the first method that we are aware of that produces approximately unbiased estimates of the genetic component of population differences in phenotype (%*G*). In particular, estimating %*G* using only genome-wide significant associated SNPs is expected to produce downward biased estimates. Indeed, when we estimated %*G* for height along PC1 in UK Biobank data using a set of 1,131 genome-wide significant SNPs (P < 5 x 10^−8^; LD-pruned to *r*^2^<0.01^26^), we obtained an estimate of 27.6% (s.e.=0.1% using s.e. of effect size estimates of each SNP, which does not account for effects of genetic drift; s.e.=7.2% using block-jackknife, which account for drift but may not be valid for small sets of SNPs), which is much lower than the estimate produced by PopDiff. When we estimated %*G* for red hair pigmentation along PC1 using a set of 47 genome-wide significant SNPs (P < 5 x 10^−8^; LD-pruned to *r*^2^<0.01), we obtained an estimate of 5.6% (s.e.=0.1% using s.e. of effect size estimates of each SNP; s.e.=8.2% using block-jackknife). More generally, there exist several methods that shrink estimated effect sizes for the purpose of maximizing polygenic prediction accuracy^3,32-35^, but these approaches are also expected to produce downward biased estimates of %*G*.

Our work has several limitations. First, all methods for detecting polygenic section may produce false-positive signals if association statistics are confounded by uncorrected population stratification^28,29^, thus careful consideration of possible stratification is required. Our secondary analyses involving homogenous subsets of samples suggest that our results are robust to population stratification (Table S14). Second, the approach for LD matrix regularization^24^ employed by PopDiff introduces a bias-variance tradeoff in estimates of %*G* (Figure 1). Estimates of %*G* may not be perfectly unbiased, although they are close to unbiased in our simulations across a broad set of genetic architectures. However, the regularization parameter that optimizes this bias-variance tradeoff may vary across different data sets (e.g. depending on the sample size and SNP set), such that analyses of new data sets may require revisiting the choice of regularization parameter; investigating other LD matrix regularization approaches may also prove useful^36,37^.

Third, we restricted our analyses to genotyped SNPs (due to complexities of LD matrix regularization and to computational cost); analyses of the %*G* explained by a larger set of genotyped and imputed SNPs might yield slightly larger estimates, consistent with the slightly larger heritability that they explain^38^. Fourth, we focused on British-ancestry samples in UK Biobank, which have very subtle structure. We did not apply PopDiff to estimate continental-level population differences in phenotype, because the much larger amount of drift between continental populations (and possible effects of differential LD^39^) will lead to large jackknife s.e., limiting power to detect polygenic section. Fifth, when polygenic selection is detected, we are unable to infer when the selection occurred, as the population structure of British-ancestry samples may often reflect differing proportions of ancestry from more deeply diverged source populations in which selection might have occurred^13,20^. Despite these limitations, PopDiff is a powerful method for quantifying the genetic component of population differences in phenotype to detect signals of polygenic selection.

## URLs

Software implementing the PopDiff method will be released prior to publication as a publicly available, open-source software package at https://www.hsph.harvard.edu/alkes-price/software; UK Biobank www site, http://www.ukbiobank.ac.uk/; LDSC software, https://github.com/bulik/ldsc/; PLINK2.0, https://www.cog-genomics.org/plink/2.0/; EIGENSOFT, https://www.hsph.harvard.edu/alkes-price/software.

## Acknowledgements

We are grateful to A. Dahl, M. Sohail, R. Maier, D. Reich and S. Sunyaev for helpful discussions. This research was conducted using the UK Biobank Resource under Application #14292 and was funded by NIH grants R01 HG006399 and R03 ES027902.

## Methods

### PopDiff method

The PopDiff method estimates the genetic component Δ*G* of population differentiation Δ*Y* (defined as the correlation between continuous ancestry gradient and phenotype *Y)*, where “genetic component” refers to the component explained by a specified set of SNPs. A genetic component that is significantly different from 0 after accounting for effects of genetic drift is indicative of polygenic selection.

We assume a simple linear model,

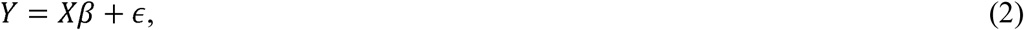

where *Y* is standardized phenotype of *N* samples, *X* is the standardized genotype (N individuals x M SNPs), β is the vector of causal effect sizes, and *?* is noise.

Let *η* denote a standardized continuous gradient of genetic ancestry, *e.g.* birth coordinates or values of a top PC (estimated in a finite sample). The population differentiation Δ*Y* and its genetic component Δ*G* are defined as follows:

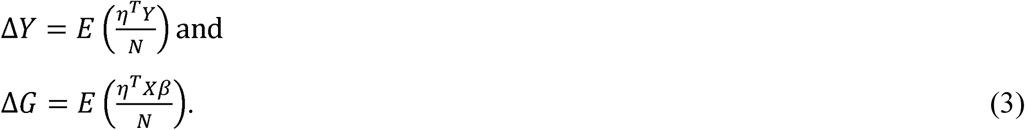

The genetic proportion of population differentiation (%*G*) is estimated as 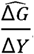.

Letting 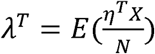denote SNP loadings along the genetic ancestry gradient *rη* it follows that

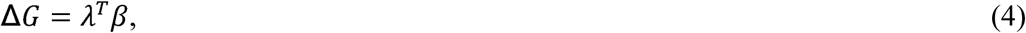

so that Δ*G* can be estimated using estimated causal effect sizes and SNP loadings.

Given marginal effect size estimates 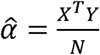 and an LD matrix estimated as 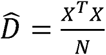, an unbiased estimate of causal effect sizes can be computed as:

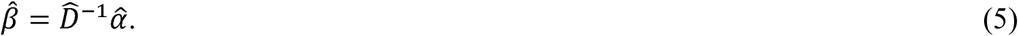

To improve computational efficiency, we divide the genome into non-overlapping blocks of 10,000 SNPs. To reduce noise in LD estimates for SNPs that are far apart, we band the LD matrices to bands of 200 SNPs, with LD estimates outside the bands set to zero. We compute local banded LD matrices 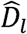 for each block. To further reduce estimation noise, we regularize LD estimates as previously described^24^ (using a regularization parameter γ), such that regularized causal effect size estimates for each block are computed as:

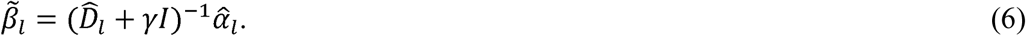

Regularized causal effect size estimates 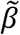 are computed genome-wide and the genetic component is estimated as

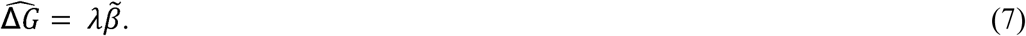

Standard errors of 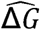 are estimated via block-ackknife, partitioning the genome into 200 blocks of non-overlapping SNPs.

We note that if the ancestry gradient *η* is a genetic PC (e.g. PC1), and summary association statistics 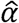 are computed by including genetic PCs as covariates, then this choice of 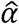 would be mathematically guaranteed to produce a 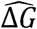 estimate of 0 along *η.* In detail, the singular value decomposition of *X* = *UYV^T^.* When *η* is PC1, 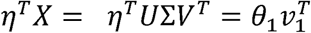, where θ_*i*_ is the i^th^ diagonal entry of Σ, and 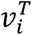 is the i^th^ row of *V*^T^. Thus, 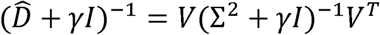. After correcting for the top 10 PCs, 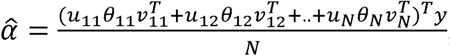, where *u*_*i*_ is the i^th^ column of *U* and y is phenotype *Y* corrected for 10 PCs. Following Equation 1, 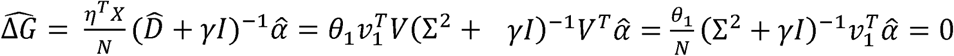.

To overcome the problem of 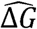 estimates being mathematically guaranteed to equal 0, when applying PopDiff to genetic PCs we partition the complete set of samples into 10 random non-overlapping subsets, compute summary association statistics for each subset using PCs of each subset as covariates, compute 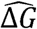 estimates using Equation 1, where η is the genetic PC of the complete set of samples and averaged 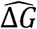 estimates across subsets. Our simulations showed that this approach produces close to unbiased 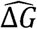 estimates (Figure 1). In the UK Biobank data that we analyzed, the average correlation between SNP loadings for PC1 of random subsets and PC1 of the entire set of 337,536 unrelated British-ancestry samples was equal to 0.93.

In secondary analyses (see Population stratification section), we instead partitioned the complete set of samples into 10 non-overlapping subsets ordered by PC1 values (similar values of PC1 within each subset, so as to minimize stratification). In the UK Biobank data that we analyzed, the average correlation between SNP loadings for PC1 of these subsets and PC1 of the entire set of 337,536 unrelated British-ancestry samples was equal to 0.027, confirming that stratifying samples by PC1 values largely eliminates stratification along PC1 (although this may not eliminate all population stratification).

### UK Biobank data set

The UK Biobank data set contains 805,426 genotyped SNPs and 488,377 samples. We removed SNPs that were multi-allelic, had a genotyping rate less than 99%, had a minor allele frequency (MAF) less than 1%, or were not in Hardy-Weinberg equilibrium (p < 10^−6^). We removed samples of non-British ancestry, samples with a genotyping rate less than 98% were removed, and related samples. After these QC filters, 516,086 SNPs and 337,536 samples remained.

When computing PCs, we LD-pruned the set of SNPs to *r*^2^<0.2 and removed regions of long-range LD and regions with significant or suggestive selection signals, as previously described. We computed PCs using the FastPCA software implemented in EIGENSOFT (see URLs).

### Simulations

We performed simulations to evaluate the bias, type I error and power of Popdiff, using real UK Biobank genotypes (*M*=516,086 SNPs). Phenotypes were simulated using an additive model

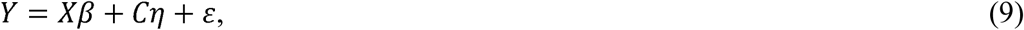

where *Xβ* represents genetic effects (including genetic effects that are correlated to ancestry) and *Cη* + *ε* (corresponding to in Equation 2) represents environmental (including environmental effects that are correlated to ancestry: *C*η). Thus, in these simulations, population differentiation Δ*Y* may be due to genetic and/or environmental effects. We specified *η* using either PC1 (computed using 337,536 British samples) or North-South birth coordinates. We simulated phenotypes for either 33,754 randomly selected samples (PC1 simulations) or all 337,536 British samples (North-South birth coordinate simulations), consistent with our analyses of real traits. Causal effect sizes β were specified with the proportion of causal SNPs *(p*) set to 1%, 10%, or 100%, via a point-normal distribution: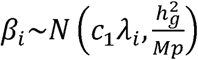 with probability *p* and 0 otherwise, where λ_i_ is the SNP loading of SNP *i* along *η*, 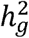 was set to 0.2, and *M* is the number of SNPs. In null simulations, *c*_*1*_ was set to 0, so that population differentiation Δ*Y* was entirely non genetic. In causal simulations, *c*_*x*_ was set to values such that *c*_*1*_*p* = 0.006. Values of Δ*Y* Δ*G* and %*G* for each simulation are provided in the Results section. Marginal effect sizes 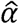 were estimated by linear regression using the top three in-sample PCs as covariates. LD matrices 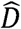 were computed using the complete set of UK Biobank British samples (*N*=337,536, *M*=516,086). *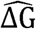* estimates were computed using Equation 1. For each type of simulation, we performed 1,000 simulations using PC1 (33,754 samples simulated) and 100 simulations using North-South birth coordinates (337,536 samples simulated).

### Analyses of UK Biobank traits

We considered 43 UK Biobank traits, restricting to 337,536 unrelated British-ancestry samples (average *N=* 321,389 phenotyped samples; Table S10). We estimated genetic components of population differentiation along three continuous gradients of genetic ancestry: PC1, North-South birth coordinate, and East-West birth coordinate (correlations between these ancestry gradients are reported in Table S24). In all analyses, banded LD matrices were computed using the complete set of British samples (N=337,536 and M=516,086). In the PC1 analysis, we divided the British samples into 10 random non overlapping subsets (6 subsets with *N*=33,754 and 4 subsets with *N*=33,753). We computed summary association statistics for each of the 10 random subsets using PLINK 2.0 (see URLs) and included the top 10 PCs of each subset, age, sex, genotyping array and assessment center as covariates. In the analysis of North-South and East-West birth coordinates, we computed summary association statistics of the complete set of British samples (N=337,536) using PLINK 2.0 and included the top 10 PCs, age, sex, genotyping array and assessment center as covariates. 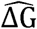 estimates were computed using Equation 1.

